# Ion channel depolarization increases repulsions between positive S4 charges to drive activation

**DOI:** 10.1101/691881

**Authors:** H. R. Leuchtag

## Abstract

The positively charged residues, arginine and lysine, of the S4 segments of voltage-sensitive ion channels repel each other with Coulomb forces inversely proportional to the mean channel dielectric permittivity ε. Dipole moments induced at rest potential in the branched sidechains of leucine, isoleucine and valine lend high values of ε to the channel. High ε keeps electrostatic forces small at rest, leaving the channel in a compact conformation closed to ion conduction. On membrane depolarization beyond threshold, the repulsive forces between positive S4 charges increase greatly on a sharp decrease in ε due to the collapse of induced dipoles, causing an expansion of the S4 segments, which drives the channel into activation. Model calculations based on α helical S4 geometry, neglecting the small number of negative charges, provide estimates of electrostatic energy for different values of open-channel ε and numbers of positive S4 charges. When the *Shaker* K^+^ channel is depolarized, the repulsion energy in each S4 segment increases from about 0.2 kcal/mol to about 120 kJ/mol (30 kcal/mol). The S4 expansions lengthen and widen the pore domain, expanding the hydrogen bonds of its α helices, thus providing sites for permeant ions. Ion percolation via these sites produces the stochastic ion currents observed in activated channels. The model proposed, Channel Activation by Electrostatic Repulsion (CAbER), explains observed features of voltage-sensitive channel behavior and offers predictions that can be tested by experiment.

**SIGNIFICANCE STATEMENT:** Science walks on two legs, experiment and theory. Experiment provides the facts that theory seeks to explain; the predictions of a theoretical model are then tested in the laboratory.

Rigid adherence to an inadequate model can lead to stagnation of a field.

The way in which a protein molecule straddling a lipid membrane in a nerve or muscle fiber responds to a voltage change by allowing certain ions to cross it is currently modeled by simple devices such as gated pores, screws and paddles. Since molecules and everyday objects are worlds apart, these devices don’t provide productive models of the way a voltage-sensitive ion channel is activated when the voltage across the resting membrane is eliminated in a nerve impulse. A change of paradigm is needed.

Like all matter, ion channels obey the laws of physics. One such law says that positive charges repel other positive charges. Since each of these ion channels has four “voltage sensors” studded with positive charges, they store repulsion energy in a membrane poised to conduct an impulse. To see how that stored energy is released in activation, we must turn to condensed-state physics. Recent advances in materials called ferroelectric liquid crystals, with structures resembling those of voltage-sensitive ion channels, provide a bridge between physics and biology. This bridge leads to a new model, Channel Activation by Electrostatic Repulsion,

Three amino acids scattered throughout the molecules have side chains split at their ends, which makes them highly sensitive to changing electric fields. The calculations that form the core of this report examine the effect of these branched-chain amino acids on the repulsions between the positive charges in the voltage sensors. The numbers tell us that the voltage sensors expand on activation, popping the ion channel into a porous structure through which specific ions are able to cross the membrane and so carry the nerve impulse along.

This model may someday enable us to learn more about diseases caused by mutations in voltage-sensitive ion channels. But for now, the ball is in the court of the experimentalists to test whether the predictions of this model are confirmed in the laboratory.

## INTRODUCTION

The voltage-sensitive or voltage-gated ion channel is a member of a superfamily of signal-transduction proteins intrinsic to biological membranes (1, 2). These glycoprotein macromolecules, embedded in the phospholipid bilayer separating cytoplasmic and external aqueous media, carry out physiological functions including the transfer of information along nerve and muscle fibers and at axon terminals. Voltage-sensitive ion channels (VSICs) include monomeric sodium (Nav) and calcium (Cav) channels and tetrameric potassium (Kv) channels. Mutations in VSICs cause diseases of brain, muscle and other organs (3).

This paper deals with membrane activation, the onset of stochastic currents of permeant ions in response to threshold depolarization of the membrane from its resting potential, negative relative to the extracellular solution, with a mean electric field of ∼14-18 MV/m. Although much experimental data is available, an explanation from first principles of the activation of these macromolecules remains to be developed. The goals of this paper are to critically examine the bases of currently accepted models; to show that these models are inappropriate at the molecular scale and contrary to published data; to propose an alternative model based on condensed state physics; to calculate certain quantitative predictions of this model, and to apply these to further our understanding of the electromechanical process of channel activation.

A fundamental question that remains to be answered is: *How does a critical reduction in the magnitude of the membrane voltage from its resting potential lead to the appearance of stochastic currents of selected ions?* We will first examine currently accepted models critically and show their inadequacy to answer this question, thereby demonstrating the need for new paradigm.

### Can an ion channel possess rigidly moving components?

Certain mechanistic models based on macroscopic devices have been proposed to explain VSIC function. The *gated-pore* model holds that VSICs are pierced by a water-filled pore fitted with a voltage-controlled gate that regulates the ion current (1,4). A variant of the gated-pore model is the *helical-screw* model, which proposes an α-helical S4 segment as rigidly advancing outward and rotating through the surrounding channel substance (5, 6). In a model based on data from a crystallized bacterial potassium channel, KvAP, certain segments are postulated to form a *paddle* that moves under voltage control within the lipid bilayer (7, 8).

While the experimental results motivated by models based on devices with rigidly moving parts have provided valuable information, their interpretation by these models cannot provide robust explanations of channel function. The tacit assumption that the S4 segment moves outward through the channel as a rigid body ignores the fact that the loops of the α helix are interconnected by hydrogen bonds, which are weak compared to interactions of the S4 segment with neighboring segments and, as shown in the calculations below, to the mutual repulsion energies of its positive charges in a depolarized channel. The properties of matter at the molecular level are vastly different from properties such as rigidity that characterize macroscopic matter. Therefore, devices with rigidly moving parts are *incommensurate with the molecular scale* and must be abandoned as models of ion-channel function.

Nonrigid motions of S4 segments or flexible structures within them were proposed by Guy and Conti (9), Durell and Guy (10) and Aggarwal and MacKinnon (11), but encountered the objection that the energy cost of breaking the segment’s hydrogen bonds is tens of kcal/mol, taken to be too high to make this concept realistic (12). However, the calculations below show that the increase in repulsion energy on activation is sufficient to account for an electrostatically driven extension of the S4 segments.

The gated-pore model and its variants are also inconsistent with single-channel data: Although the current response observed in an axon membrane with mm-size electrodes follows the field smoothly, the response observed with a μm-size patch clamp to record currents from the nm-size channel is a stochastic sequence of unit currents (13). In contrast to the predictable effect of opening a macroscopic gate, a channel opening thus only predicts the *probability* of onset and termination of unit currents (14). The ion channel is not a simple gated pore.

To explain channel function from first principles, such device-based models must be abandoned and replaced in a paradigm shift by an approach based on molecular concepts.

### Electrostatic interactions between positive charges in an S4 segment

The four S4 segments of all VSICs are highly ordered structures, and that order has been strongly conserved in evolution. As a common motif, each possesses an array of *positively charged* amino acid residues, arginine (R) and lysine (K), separated by (predominantly) two uncharged, nonpolar residues. The S4 segments are recognized as the components that respond to changes in voltage across the membrane; they are part of the *voltage sensor domain* (segments S1-S4) of VSICs (4, 15). The fact that electric charges repel as well as attract is well known, and its implications for the biophysics of ion channels, pointed out by Leuchtag (16) are here applied to help explain channel activation.

The claim that the positive S4 charges are neutralized by salt bridges with negatively charged residues (17, 18, 19) and hence do not repel each other, must be rejected in view of the facts that the positive residues greatly outnumber the negative ones, that the positively charged residues are conserved and that they play an important role in channel activation (10). The finding by Stühmer et al. (20) that a reduction of total equivalent positive charge can reduce voltage dependence in neutralization mutants of a sodium channel implies that the mutual repulsion of the positive S4 charges cannot be ignored.

The present paper presents a computation from the laws of electrostatics of the repulsion energies between the positively charged residues in a simple model of the S4 segments. An electromechanistic model, Channel Activation by Electrostatic Repulsion (CAbER), is proposed to explain the role that these repulsive forces play in ion-channel activation; a preliminary report has been presented (21).

## METHODS

### Electrostatics of an S4 segment

The electrostatic interaction is a long-range force between charged particles in a nonconducting medium. Coulomb’s law for charges *q*_*i*_ and *q*_*j*_ separated by distance *r*_*ij*_ with unit vector **u**_ij_ between them in a medium of dielectric permittivity ε, in MKS units, is

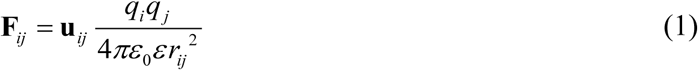

where ε_0_ = 8.854 × 10^-12^ C^2^/Nm^2^ (22).

After a step depolarization is imposed on a VSIC, it is no longer in equilibrium. Its response is a transient process during which charges move. Since ion-channel structures must move during activation, electrostatic calculations based on equations that assume equilibrium conditions, such as the Poisson–Boltzmann equation (23), are not applicable to channel activation.

In considering the way in which the external electric field must move charges, Hille (24), finds that elimination of certain alternative hypotheses leaves only a *direct action* on charges that are part of the channel, as earlier suggested by Hodgkin and Huxley (25). The charges acted upon were taken to be the positive charges on the S4 segments (26, 27, 28, 29). However, outward S4 motions are observed for depolarizations less in magnitude than the resting potential, which leave the inner membrane surface negative and continuing to attract the positive S4 charges inward, not outward as the direct action hypothesis assumes. Since the membrane voltage acts across the entire membrane and is not in series with a conducting element, subtraction of the resting potential from the membrane voltage is not allowed (30). Therefore, the process responsible for the outward movement cannot be a *direct* response to the change in membrane voltage.

Rather, the activation process must involve an *intermediate variable* that reacts to the voltage change and in turn leads to the observed outward motions of the positive charges. Since by Eq. 1 the only variable dependent on the channel environment is the dielectric permittivity ε, that intermediate variable must be ε.

Capacitive currents called “gating currents” have been measured in excitable membranes when ionic currents are blocked (31). Since the dielectric permittivity is the dimensionless parameter of capacitance, the gating currents must be seen as indications of the changes of ε in the VSICs during activation. To further pursue the effects of a variable ε, let us consider the classical electrodiffusion model.

### Ferroelectricity in biological structures

The classical *electrodiffusion model* of excitable membrane function was found to be inadequate because of its inability to fit ion-channel data such as the steep rectifications observed in squid axon (32). However, the rejection of electrodiffusion *in general* was premature, as the classical model is based only on calculations in which the parameters of the Nernst–Planck equation, ion mobility and dielectric “constant” ε, are *held constant* (33). Following the model of Hodgkin and Huxley (25), which postulated voltage-dependent conductances, Leuchtag (34) took ε to be a function of *E*, the magnitude of the electric field **E** across the membrane. Under the hypothesis that ε = ε(*E*), the electrodiffusion equations become identical to those that describe *ferroelectric* materials (35, 36).

Ferroelectrics have built-in electric fields. Ferroelectric materials can sense temperature changes (pyroelectricity), interchange electrical and mechanical effects (piezoelectricity) and produce electro-optic effects (light scattering and field-sensitive birefringence). By definition, a ferroelectric phase is characterized by a nonzero electric polarization (dipole moment per unit volume) that is reversible by an external field. (However, since no field reversal occurs in activation, this defining property is not a feature of the CAbER model.) The *spontaneous polarization* of ferroelectrics exists between definite temperature limits, the upper and lower Curie points. In the CAbER model, the *field effect*, in which the (upper) Curie point rises with increasing external electric field, is critical to the explanation of ion-channel activation.

### Excitable membranes as ferroelectric liquid crystals

Biological membranes are neither crystalline solids nor disordered liquids; rather, they are phases with order intermediate to these, liquid crystals or mesophases (37). Although bulk liquid crystals are usually classified as nematic, cholesteric and smectic phases, these categories are inadequate to cover the complexities of ion channels in excitable membranes. *Blue phases*, ferroelectric materials characterized by a double twist such as two helices in contact at an angle, exist in narrow temperature ranges with cubic or amorphous (fog) symmetries (38).

Excitable membranes share a number of similarities to ferroelectric liquid crystals, including transition temperatures known as heat block and cold block in axons, surface charges, temperature-stimulated currents, and both thermal and current–voltage hysteresis (33, 39). Light scattering (40) and voltage-sensitive birefringence, typical of ferroelectric liquid crystals (41), were observed in excitable membranes (42).

Leuchtag (43) fitted the ferroelectric Curie-Weiss law with a Curie temperature of 49.8 °C to data by Palti and Adelman (44) on the capacitance of squid axon as a function of temperature. Cole–Cole plots of frequency-domain data from sodium channels in squid axon membrane (45) are semicircles similar to those measured by Wróbel et al. (46) in ferroelectric liquid crystals, in which the permittivities were strongly bias field dependent. The hypothesis of ferroelectric phases in VSICs (33, 34, 47, 36, 39) was extended to soliton propagation and magnetic effects in axons (48).

Liu et al. (49) observed ferroelectric behavior, including polarization–field hysteresis, butterfly-shaped deformation–voltage loops and voltage-sensitive switching of polarization, in dried intimas of porcine aortic walls. Evidence of ferroelectricity in peptide nanotubes (50) supports the concept of a ferroelectric phase in these biomimetic structures. Ermolina et al. (51) observed liquid-crystal-like ferroelectric behavior in bacteriorhodopsin, an integral protein of the purple membrane of *Halobacterium salinarium.*

### Voltage-sensitive ion channels as bioferroelectric liquid crystals

Ferroelectric phases exist in crystals (35), polymers (52), liquid crystals (53) and submicrometer films (54). The related *antiferroelectric* phases (41) have alternating opposite tilt angles, and since the α helical segments are macrodipoles (55, 56) that thread back and forth across the membrane, the channel may have antiferroelectric properties. Ferroelectric liquid crystals respond to threshold changes in external electric and magnetic fields (Frederiks’s effect) and to bending and torsion (flexoelectric effect) (57). The dielectric permittivity of a ferroelectric liquid crystal is strongly dependent on the electric field (46).

Table 1 shows that the transmembrane segments of voltage sensors in the four repeats of VSICs have structural features characteristic of ferroelectric and antiferroelectric liquid crystal molecules: chiral centers, resonance-stabilized aromatic rings, C=O dipoles and aliphatic chains (58).

**Table 1.**
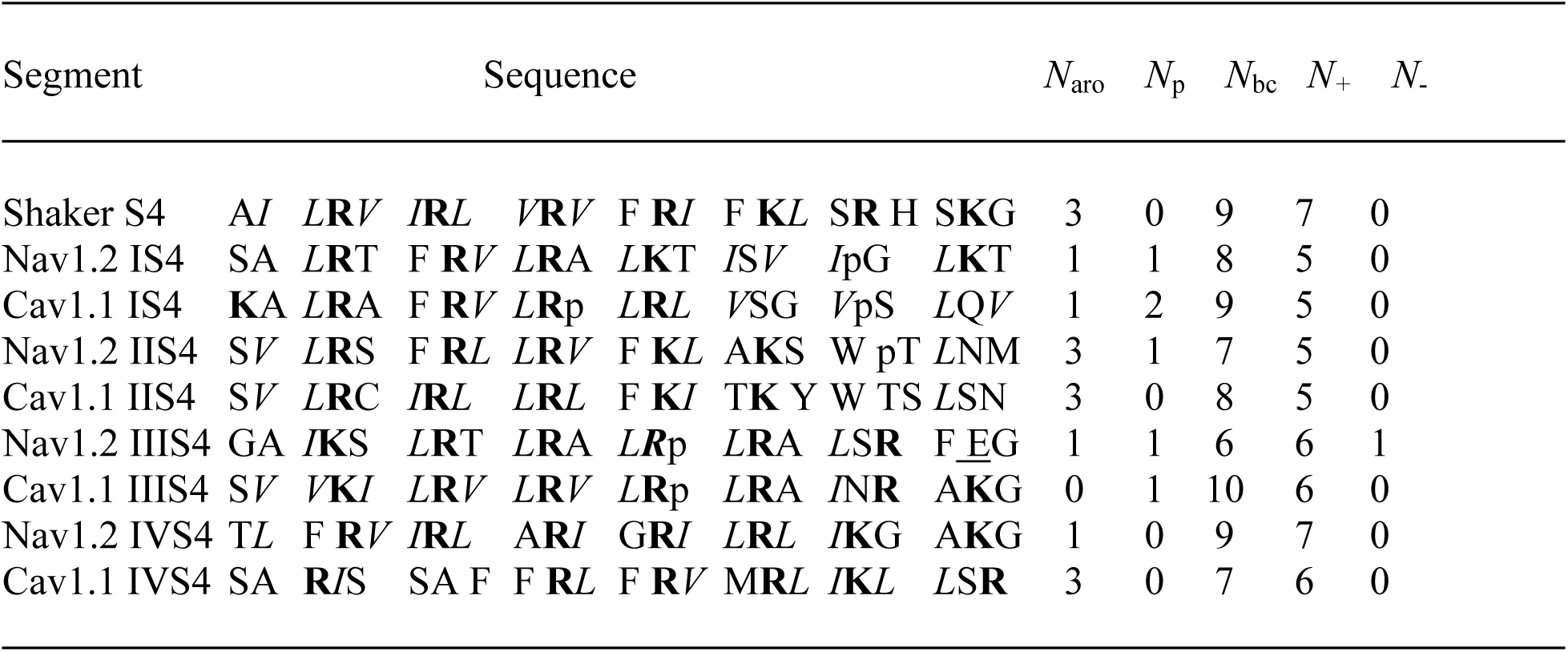
Structural characteristics in homologous sequences of S4 segments for selected channels, adapted from Hille (1), p. 607. *N*_aro_ = number of aromatic residues (F, Y, W and H) in the S4 segment *N*_p_ *=* number of prolines (p) *N*_bc_ = number of branched-chain *I, L* and *V* residues *N*_+_ = number of positively charged **R** and **K** residues *N*_-_ = number of negatively charged <u>D</u> and <u>E</u> residues.

## THE BIOFERROELECTRIC HYPOTHESIS

Since the ion channel has properties similar to ferroelectric phases but is structurally different from bulk ferroelectrics, it is referred to as *bioferroelectric* (59, 36, 60, 39). The CAbER hypothesis proposes that the bioferroelectricity of the excitable membrane originates in the α helices of the VSIC. This is supported by the parallels between α helices and ferroelectric materials, such as hydrogen bonding and domain formation, reminiscent of antiparallel α-helical bundles, as pointed out by Gilson and Honig (23). An implication of dipole–charge interactions, they continue, is that long-range interactions may play important roles in protein function.

A potential difficulty with the bioferroelectric hypothesis is that a ferroelectric phase must exceed a certain minimum size (39). This issue was clarified when Bune et al. (54) showed that polymer films of thickness 1.0 nm, less than the dimensions of a VSIC, can support a ferroelectric phase, indicating that this phase can propagate by coupling exclusively within the plane of the film.

The data cited above suggest that threshold depolarization lowers the Curie point to the channel temperature by the field effect, leading to a bioferroelectric phase transition. The CAbER model postulates that a critical reduction in the magnitude of the membrane electric field converts the channel from a *ferroelectric phase at rest potential* to a *nonpolar phase on threshold depolarization*. This order–disorder phase transition reduces ε, increasing the mutual repulsions of between the positive charges in the S4 segments, leading to an open channel conformation that allows conduction of permeant ions.

### Branched-chain amino acids affect channel dielectric permittivity

The S4 segments of all members of the S4-containing superfamily (61) are characterized by an array of positively charged amino acid residues, arginine (**R**) and lysine (**K**) at every third position within the segment, to form the pattern x(**R**/**K**)x x(**R**/**K**)x x(**R**/**K**)x …. (See Table 1). The positively charged residues are essential to channel function; their substitution by neutral residues alters channel activation (20, 62). The residues labeled “x” between the positively charged residues are:

- the aromatic residues, tryptophan (W), phenylalanine (F), tyrosine (Y) and histidine (H);
- frequently, a proline (p) that bends the α helical segment; and
- predominantly, the branched-chain amino acids (BCAAs) isoleucine (*I*), leucine (*L*) and valine (*V*).

The BCAAs are distributed throughout the channel. Table 1 lists the number of aromatic residues, prolines, BCAAs and positively and negatively charged residues in selected S4 segments. Negatively charged residues, aspartic acid (<u>D</u>) or glutamic acid (<u>E</u>), are seldom found in S4, but note an <u>E</u> in Nav1.2 IIIS4 in Table 1. These negative charges, as well as the positive charge of H, are often ignored for simplicity. The attraction of positive charges in S4 by the few negative charges in segments S2 and S3 (salt bridges) presents a relatively minor effect on S4 free energy, and is neglected in these simple calculations.

### Liquid crystals with branched sidechains exhibit strong ferroelectricity

Liquid crystal molecules with aromatic rings and sidechains of BCAAs have been found to exhibit ferroelectric properties with extremely large values of spontaneous polarization and dielectric permittivity (58, 63). These properties are due to their large induced dipole moment. One example is 4’-3M2CPOOB, with two benzene rings and a terminal isoleucine sidechain:

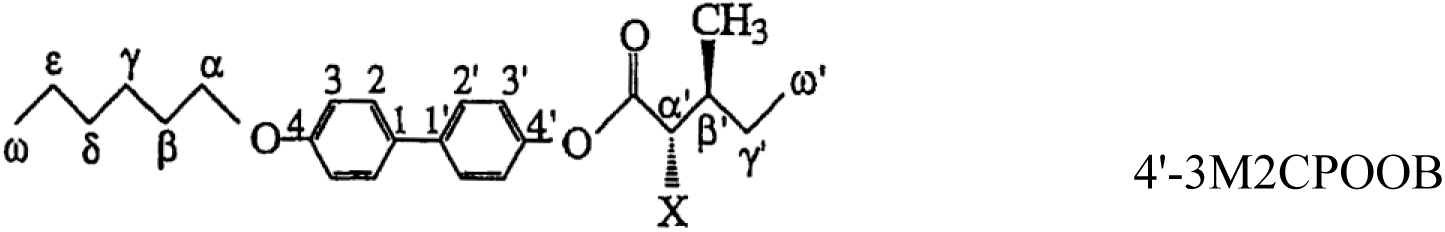

In the diagram, the α’ and β’ carbons are chiral; X is chlorine. In the ferroelectric phase of 25 μm samples, with a field strength of 500 V/cm at a frequency of 30 Hz, dielectric permittivities of over 3000 were recorded, along with a strong field effect (58).

## CALCULATIONS

### Function of branched-chain amino acids play in channel activation

VSIC segments are polymers with short sidechains on an α-helical backbone, placing them in the category of *sidechain polymers*. Many sidechain polymers similar to VSIC segments in possessing aromatic mesogens, chiral centers and branched structures exhibit ferroelectricity (52). The importance of these structures to VSIC function is demonstrated by mutation experiments.

Mutations of branched to unbranched (or even different branched) residues affects channel function: Lopez et al. (64) showed that *L* to A mutations in segment S4 strongly shift the conductance–voltage curves in the *Shaker* K^+^ potassium channel, mostly in the direction of higher voltage. Amino-acid substitution of unbranched for branched residues in fragments of electric eel Na^+^ channels inserted into bilayers affected their voltage response, indicating a link between the branched sidechains and the gating process (65). Mutations of *L* to A in S4 segments of domains I and II of human skeletal muscle Nav1.4 channels expressed in human embryonic kidney cells altered steady-state activation curves and slowed inactivation (66). Voltage-sensor motion in Ci-VSP is tuned over two orders of magnitude or shifted 120 mV by the size of an isoleucine, *I*126, of the S1 helix, which is strictly conserved over a number of K^+^ and Na^+^ channels (67). A triple mutation of uncharged residues, two of which change one BCAA into a different one, converts the kinetic and voltage-dependent properties of *Shaw* potassium channels into *Shaker*, and show that the gating currents precede the ionic currents; one of these, *I*372*L*, is primarily responsible for changes in cooperativity and voltage dependence (68).

These experimental findings show that BCAAs play an important role in channel activation, a role that can be accounted for by the effect of these residues on the dielectric permittivity of the channel: The collective effect of the induced dipole moments is to create surface polarization charges that oppose the applied field within the molecule, reducing the *electric displacement*, **D** = ε**E**, by the factor ε (22); see Figure 1.

**Figure 1.**
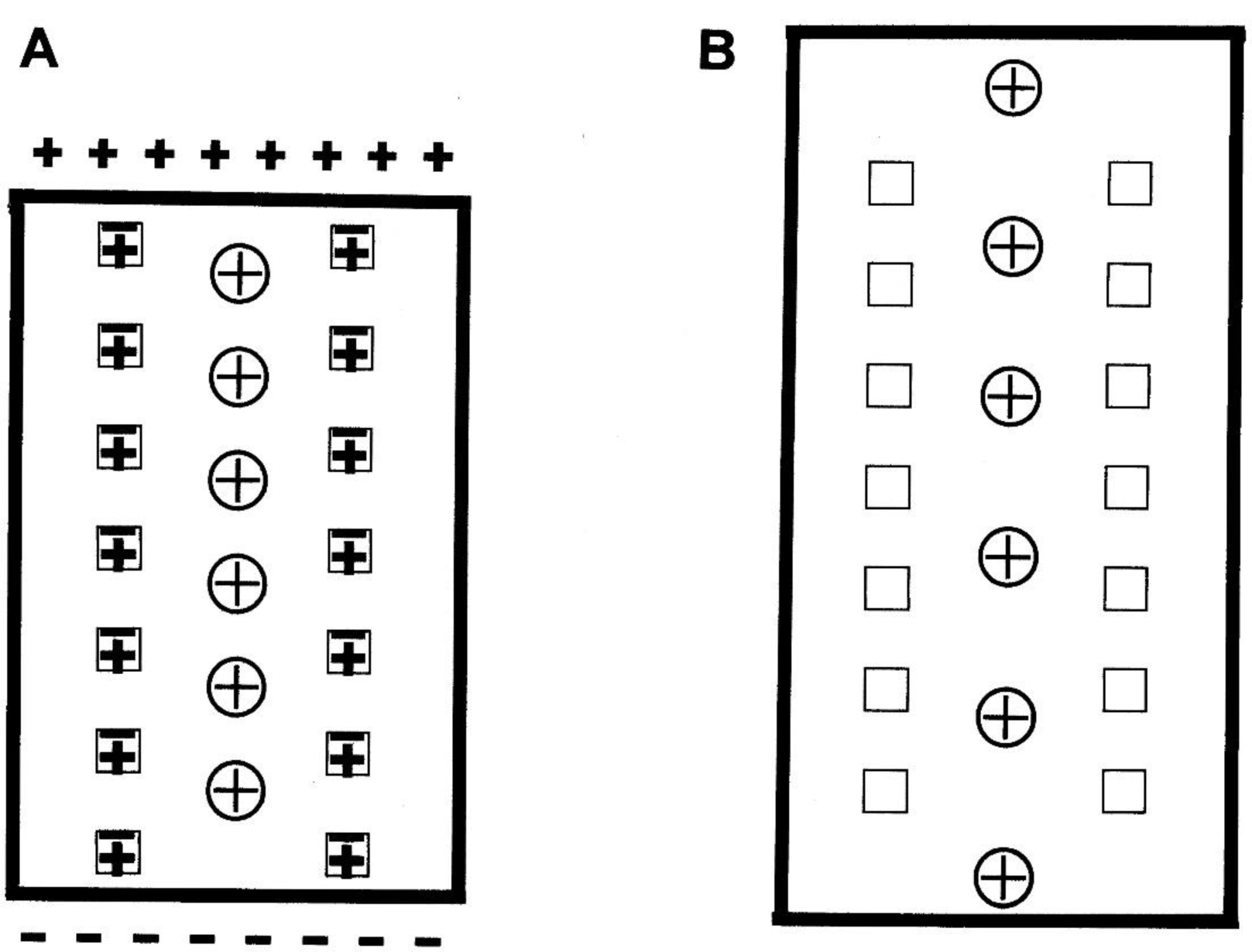
Schematic sequence of changes at channel activation in the proposed electromechanism. For simplicity, the ion channel is represented by a single column of positively charged R/K residues (+) within an S4 segment, surrounded by two columns of branched-chain amino acid I/L/V residues (squares), which are distributed throughout the channel. The outside of the channel is up. **A**. At the high electric field of the resting potential, a large dipole moment is induced in the sidechains of the BCAAs (polarized squares), partially shielding the applied electric field and thus raising the channel’s mean dielectric permittivity to a high value, say ε = 500. Because ε is in the denominator of Coulomb’s law, electrostatic forces and energies are low, and the mutual repulsions between the positive residues do not appreciably increase the helical pitch of S4 from its standard value. **B**. When the channel is depolarized beyond threshold, reducing the applied electric field, the BCAA sidechains relax and lose their dipole moments (empty squares). This reduces ε to values between 3 and 5, causing the repulsions to increase by a factor of the order of 100. As the positive residues move apart from one another, they expand the S4 segments.

### Calculation of repulsion energy within S4 segments

The S4 segment at rest potential is modeled as an α helix with its axis in the *z* direction, perpendicular to the membrane plane. The small number of negative residues and the proline bend, which occurs frequently but not always in S4 segments (see Table 1), are omitted for simplicity.

Although BCAA sidechains in the channel segments occur in larger numbers than in 4’-3M2CPOOB, in which ε rises to over 3000 in the ferroelectric phase, we will conservatively assume ε = 500 for the ferroelectric resting channel in the model calculations. In contrast, proteins in equilibrium have a dielectric constant ranging between 3 and 5 (12). We here consider only the component of ε in the transmembrane direction.

The positively charged residues are modeled as points on a cylinder of radius *R* = 0.80 nm from the helical axis (16). The charges are numbered from 1 to *N* (or *N*_+_, as in Table 1), starting with the outermost. Since the cytoplasm may be considered to be incompressible, the innermost positive charge, *q*_N_, can be considered fixed in position. When ε is so large that electrostatic forces can be neglected, the S4 segment will have the dimensions of a standard α helix, with residues on a right-handed helix of angular separation 100° and a separation in the *z* direction of 0.15 nm. Since the charges are at every third position, the axial distance between them is *A* = 0.45 nm and the angular separation θ = 300°. The distance *r*_*ij*_ between two of the positive charges, *i* and *j*, is (16):

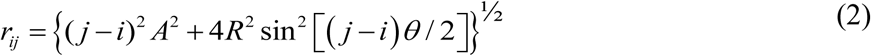

Because of the vectorial nature of force and the interactions with neighboring segments, it is simpler to calculate the electrostatic potential energy than the force. The mutual potential energy (in joules) between two charges, *q*_*i*_ = *q*_*j*_ = *e* = 1.60 × 10^-19^ C, is (22, 16)

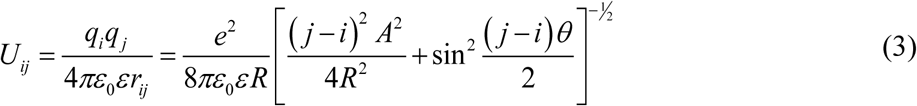

The net potential energy *U*_*1*_(ε, *N*), in kilojoules per mole, of the outermost charged residue, *i* = 1, is the sum of the energies due to the repulsion of the other *N -* 1 charges:

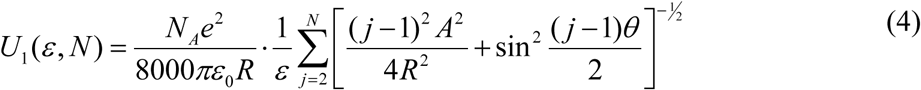

where *N*_A_ is Avogadro’s number, 6.02 × 10^23^ mol^-1^. Calculated values of *U*_*1*_*(ε, N)* in units of kJ/mol are shown in Table 2 for *N* ranging from 2 to 8, and for a resting ε of 500 and depolarized values of ε of 3, 4 and 5 (12). Calculations were carried out on a Texas Instruments TI-35X calculator and Microsoft Excel 2007.

**Table 2.** Electrostatic repulsion energy. *U*_*1*_(ε, *N*) in kJ/mol of the outermost charged residue of a model S4 segment as a function of channel dielectric permittivity ε and number *N* of positive charges; calculated from Equation 4.

| $N$ | Resting | Depolarized Channel | | |
| --- | --- | --- | --- | --- |
| | $\epsilon = 500$ | 3 | 4 | 5 |
| 2 | 0.302 | 50.3 | 37.7 | 30.2 |
| 3 | 0.470 | 78.3 | 58.7 | 47.0 |
| 4 | 0.602 | 100.3 | 75.3 | 60.2 |
| 5 | 0.724 | 120.7 | 90.5 | 72.4 |
| 6 | 0.840 | 140.0 | 105.0 | 84.0 |
| 7 | 0.943 | 157.1 | 117.8 | 94.3 |
| 8 | 1.028 | 171.3 | 128.5 | 102.8 |

## DISCUSSION

### Repulsion energy expands the S4 segment

For an S4 segment with seven positive charges, *N* = 7, as listed in Table 1 for the *Shaker K*^*+*^ channel, with a resting ε = 500, Table 2 shows an electrostatic potential energy *U*_1_ = 0.943 kJ/mol (0.225 kcal/mol), only ∼12% of the hydrogen-bond energy of ∼ 8 kJ/mol. This small perturbation is unlikely to change the widths of the backbone hydrogen bonds appreciably from the standard value. When the channel is depolarized, however, the repulsion energy increases to 157.1 kJ/mol (37.5 kcal/mol) if ε = 3, to 117.8 kJ/mol (28.1 kcal/mol) if ε = 4, and to 94.3 kJ/mol (22.5 kcal/mol) if ε = 5.

These repulsion energies on activation, more than 1100% as great as the hydrogen-bond energy, must result in the separation of the positive charges from each other and, consequently, an *expansion of the S4 segment*. Thus, the reduction of ε from a resting value of 500 to a depolarized value of 4 ± 1 results in a great increase in the repulsion energy of the outermost positive residue, which must cause the S4 segments to expand. This expansion is consistent with the observed outward movements (5, 6). Luminescence energy transfer measurements on *Shaker* K^+^ channels indicate a mean S4 displacement of 1.0 ± 0.5 nm, with a normal component of 0.5 ± 0.2 nm (69).

For an S4 segment with five positive charges, *N* = 5, as in segments IS4 and IIS4 of Nav1.2 and Cav1.1, the percentage energy perturbations change to ∼9% at rest and at least 900% when depolarized. The latter increase in repulsion energy, while substantial, is notably less than that for *N* = 7, showing that a decrease in the number of positive S4 charges reduces the effectiveness of channel activation.

The calculations show that *the sizable reduction of ε* induced by the channel’s phase transition from ferroelectric (at rest) to nonpolar (depolarized) in a threshold depolarization *leads to a sharp increase of the electrostatic repulsion* between positive residues. These repulsions must *expand the four S4 segments*.

These electrostatic calculations also show that the potential energy *U*_*1*_*(ε, N)*, as well as the S4 expansion, increases with the number of positive residues *N*. Thus S4 segments with fewer sensor charges are not as effective in channel activation as those with more. This is consistent with the experimental findings by Stühmer et al. (20) that neutralization of positive charges generally results in a decrease in sodium-channel activation, and that the decrease becomes greater as more charges are neutralized. It is also consistent with the finding by Logothetis et al. (70) that differences in the charge content of S4 segments between mammalian potassium channels, Kv1.1, and the *Drosophila Shab11* are sufficient to account for the distinct *gating valence* of each channel.

Not all seven positive charges are needed to open the *Shaker* channel. The margin of safety provided by such excess capacity may have contributed to the evolutionary pressure to increase the number of positive charges in the S4 segments of more complex organisms. The greater number of positive charges in ion channels of more advanced organisms would result in a larger repulsion force and hence a more rapid and reliable activation process, presumably providing an adaptive advantage to the organism.

Certain complications have been omitted for simplicity: the presence of a small number of negative residues in the S1 and S2 segments; the occasional irregularities in the distance between positive charges; the effects of surface charges along the aqueous boundaries and the headgroups of the lipid bilayer; the tilt angle of the S4 segments; the proline bend of many S4 segments, and the torques that cause the S4 helix to unwind as well as expand. Further studies that take these factors into consideration should modify the above calculations but are unlikely to greatly alter the major conclusions.

Papazian et al. (71) found evidence by site-directed mutagenesis and electrophysiological analysis that that the positively charged S4 residues of *Shaker* K^+^ channels are important in channel activation and inactivation, but that these are not the only amino acids that affect the channel’s voltage dependence. The increase in electrostatic repulsion during activation must drive the S4 segments apart from each other within the membrane plane (72), as well as expanding them. Weinreb and Magura (73) propose that this sideways movement reduces the force of repulsion between the S4 charges and a cation traversing the channel axially; although this model is qualitatively valid, the calculation overestimates the repulsion energy in the resting state by assuming that all the charges of an S4 segment are concentrated at a single point.

### Widening of the pore segments allows stochastic currents of permeant ions-

The S4 expansions demonstrated above result in a global expansion of the channel into a new equilibrium conformation, the open state of the VSIC. Studies of the physical chemistry of hydrogen bonds (74) show that when they are widened, hydrogen bonds become subject to occupation by ions such as K^+^, Na^+^, Li^+^ and Ca^2+^. This ion exchange permits bare ions to enter a conduction pathway formed by the four α helices of the S6 segments (4), and occupy sites within it (16, 39). Permeant ions, selected and stripped of their hydration waters, enter widened hydrogen-bond sites in the backbones of the pore segments, where they become mobile.

During transient periods when a critical ion concentration of permeants is reached in the pore domain, the ions form a connected pathway and hop cooperatively from site to site. This *percolation* (75) across the channel is driven and directed by the electrochemical potential difference between the aqueous phases. As the ion occupation pattern changes unpredictably from moment to moment due to thermal motions, the ion current is not continuous in time but broken into stochastic bursts, as observed in single-channel currents (13).

### Structure–function relations

The CAbER model describes a way in which the functional properties of channel activation arise from its structural features, including positively charged arginines and lysines, chiral carbon atoms, branched sidechains and aromatic groups. It accounts for:

- the resting potential as a necessary condition for molecular excitability;
- the role of branched-chain amino acid residues in channel activation;
- the indirect effect of membrane depolarization on channel activation;
- a quantitative relation between the number of positive charges in S4 and the electrostatic energy of channel activation; and
- a stochastic electromechanism with unpredictable onsets and terminations of ion current.

### Model predictions

No other model has been able to account for these phenomena from physical principles. The validity of the CAbER model may be tested by experiments designed to falsify or support the following predictions of the hypothesis:

- In the activation of a channel by a depolarization, the distance between any two charged residues in S4 will not remain constant but will increase if the charges are like and decrease if they are unlike.
- Replacement of one or more branched residues with unbranched ones, or vice versa, will shift the activation curve.
- Replacement of aromatic residues with nonaromatic ones, or vice versa, will affect activation.
- Channel activation exists in a temperature range with upper and lower limits, accounting for the heat block and cold block observed in axons. In wild-type channels, the values of these limits will in general vary with the type of channel, consistent with the adaptation of the organism to its environment.

## CONCLUSIONS

The conserved motifs of regular arrays of positively charged residues in the voltage sensors provide a clear sign that repulsive forces play significant roles in ion channel activation. A threshold voltage reduction leads to a cooperative relaxation of the sidechains of the branched chain amino acids. This reduces their dipole moments and changes the phase of the channel from bioferroelectric (resting) to nonpolar (first stage of activation). The dielectric permittivity of the channel thereby drops from a high to a low value, greatly amplifying all electrostatic forces, including the repulsive forces between the positive charges in S4. These repulsions expand the four S4 segments. Because of interactions between segments, these movements affect the global conformation of the channel (second stage of activation).

Two assumptions were made to explain the basis of channel activation: (1) The mean dielectric permittivity ε is an increasing function of electric field magnitude. (2) The effect of branched nonpolar amino acids on channel dielectric permittivity in ion channels is similar to that in ferroelectric liquid crystals.

Reasoning from these assumptions shows that voltage sensitivity is not due to the direct action of the electric field on the positive charges of the S4 segments but due to an indirect effect: The high electric field across the resting channel creates induced dipole moments in the branched-chain amino acid residues, creating a bioferroelectric phase with a polarization that reduces the applied field. Thus the mean dielectric permittivity of the channel is high at rest, so that the repulsions between the positive charges are small, leaving the resting channel compact and impermeable.

On threshold depolarization, the branched sidechains relax and lose their dipole moments, greatly reducing the permittivity and so increasing the repulsive forces between the positive residues in S4. When the *Shaker* K^+^ channel is depolarized, the repulsion energy in each S4 segment increases to about 120 kJ/mol (30 kcal/mol). These repulsions cooperatively expand the S4 segments, driving a conformational change of the ion channel to an expanded configuration with sites along which permeant ions stochastically hop across the membrane until inactivation intervenes.

The energetics of this effect for the simple model used for the above calculations show that the S4 segments must expand. The calculations quantitatively describe the expansion of the S4 segments induced by the lowering of the channel’s dielectric permittivity when the branched sidechains lose their dipole moments. While certain structural complications have been omitted for simplicity, improvements are unlikely to alter the conclusion that powerful repulsions between positive charges of the S4 segments are indirectly induced by a threshold depolarization.

The molecular electromechanism approach to voltage-sensitive ion channels presented in the Channel Activation by Electrostatic Repulsion model explains existing data and offers predictions offers predictions that can be tested by experiment.

## Author Contributions

H. R. Leuchtag is the sole author of this manuscript.

## Acknowledgments

I thank Vladimir Bystrov, Hervé Duclohier, Saȉd Bendahhou, Fredrik Elinder and Peter Stiles for helpful comments.

## REFERENCES

1. Hille, B. 2001. Ion Channels of Excitable Membranes. 3rd Ed. Sunderland: Sinauer.

2. Leuchtag, H. R. 2008. Voltage Sensitive Ion Channels: Biophysics of Molecular Excitability. Dordrecht: Springer

3. Ashcroft, F. M. 1999. Ion channels and disease. Academic Press, San Diego.

4. Börjesson S. I. F, and F. Elinder. 2008. Structure, function, and modification of the voltage sensor in voltage-gated ion channels. Cell Biochemistry and Biophysics 52:149–174.

5. Catterall, W. A. 1986. Voltage-dependent gating of sodium channels. Correlating structure and function. Trends Neurosci. 9:7–10.

6. Guy, H. R., and P. Seetharamulu. 1986. Molecular model of the action potential sodium channel. Proc. Natl. Acad. Sci. USA. 83:508–512.

7. Jiang, Y., V. Ruta, J. Chen, A. Lee, and R. MacKinnon. 2003. The principle of gating charge movement in a voltage-dependent K^+^ channel. Nature 423:42–48.

8. Yarov-Yarovoy, R.V., D. Baker, and W.A. Catterall. 2006. Voltage sensor conformations in the open and closed states in rosetta structural models of K^+^ channels. Proc. Natl. Acad. Sci. USA. 103:7292–7297.

9. Guy, H. R., and F. Conti. 1990. The propagating helix model of voltage-gated channels. Biophys. J. 57:111A.

10. Durell, S. R., and H. R. Guy. 1992. Atomic scale structure and functional models of voltage-gated potassium channels. Biophys. J. 62:243–252.

11. Aggarwal, S. K., and R. MacKinnon. 1996. Contribution of the S4 segment to gating charge in the *Shaker* K^+^ channel. Neuron 16:1169–1177.

12. Ben-Tal N., A. Ben-Shaul, A. Nicholls, and B. Honig. 1996. Free-energy determinants of alpha-helix insertion into lipid bilayers. Biophys. J. 70:1803–1812.

13. Neher, E., and B. Sakmann. 1976. Single-channel currents recorded from membrane of de-enervated frog muscle fibres. Nature 260:779–802.

14. Colquhoun, D. A., and G. Hawkes. 1983. The principles of the stochastic interpretation of ion-channel mechanisms. In: Sakmann, B., E. Neher, editors. Single-Channel Recording. Plenum, New York, pp 135–175.

15. Gonzalez, C., G. F. Contreras, A. Peyser, P. Larsson, A. Neely, and R. Latorre. 2012. Voltage sensor of ion channels and enzymes. BioPhys. Rev. 4:1–15.

16. Leuchtag, H. R. 1994. Long-range interactions, voltage sensitivity, and ion conduction in S4 segments of excitable channels. Biophys. J. 66:217–224.

17. Durell, S.R., I.H. Shrivastava, and H.R. Guy. 2004. Models of the structure and voltage-gating mechanism of the Shaker K^+^ channel. Biophys. J. 87:2116–2130.

18. Freites, J.A., E.V. Schow, S.H. White, and D. J. Tobias. 2012. Microscopic origin of gating current fluctuations in a potassium channel voltage sensor. Biophys. J. 102:L44–L46.

19. Lecar, H., H.P. Larsson, and M. Grabe. 2003. Electrostatic model of S4 motion in voltage-gated ion channels. Biophys. J. 85:2854–2864.

20. Stühmer, W., F. Conti, H. Suzuki, X.D. Wang, M. Noda, N. Yahagi, H. Kubo, and S. Numa. 1989. Structural parts involved in activation and inactivation of the sodium channel. Nature 339:597–603.

21. Leuchtag, H. R. 2018. Eliminating warrantless assumptions facilitates consideration of an electrostatic model of ion-channel activation. 2018 Biophysical Society Meeting Abstracts Biophys. J. 114(3):480a. DOI:https://doi.org/10.1016/j.bpj.2017.11.2638

22. Jackson, J.D. 1962. Classical Electrodynamics. John Wiley, New York.

23. Gilson, M.K., and B.H. Honig. 1988. Energetics of Charge–Charge Interactions in Proteins. Proteins: Structure, Function, and Genetics 3:32–52.

24. Ref. 1, page 56.

25. Hodgkin, A. L., A. Huxley. 1952. A quantitative description of membrane current and its application to conduction and excitation in nerve. J. Physiol. (London) 117:500–544.

26. Yang, N., and R. Horn. 1995. Evidence for voltage-dependent S4 movement in sodium channels. Neuron 15:213–218.

27. Larsson, H. P., O. S. Baker, D. S. Dhillon, and E. Y. Isacoff. 1996. Transmembrane movement of the *Shaker* channel S4. Neuron 16:387–397.

28. Baker, O. S., H. P. Larsson, L. M. Mannuzzu, E. Y. Isacoff. 1998. Three transmembrane conformations and sequence-dependent displacement of the S4 domain in shaker K+ channel gating. Neuron 20:1283–1294.

29. Cha, A., and F. Bezanilla. 1997. Characterizing voltage-dependent conformational changes in the Shaker K^+^ channel with fluorescence. Neuron 19:1127–1140.

30. Leuchtag, H. R., What’s wrong with the Hodgkin–Huxley model? An exercise in critical thinking 2016 Biophysical Society Meeting Abstracts Biophys. J. 112(3):464a (2016). DOI: http://dx.doi.org/10.1016/j.bpj.2016.11.2488.

31. Armstrong, C. M, and F. Bezanilla. 1973. Currents related to the gating particles of the sodium channels. Nature 242:459–461.

32. Cole, K. S. 1965. Electrodiffusion models for the membrane of squid giant axon. Physiol. Rev. 45:340–379.

33. Leuchtag, H. R. 1987a. Indications of the existence of ferroelectric units in excitable-membrane channels. J. Theor. Biol. 127:321–340.

34. Leuchtag, H. R. 1987b. Phase transitions and ion currents in a model ferroelectric channel unit. J. Theor. Biol. 127:341–359.

35. Lines, M. E, A. M. Glass. 1977. Principles and Applications of Ferroelectrics and Related Materials. Clarendon Press, Oxford.

36. Leuchtag, H. R., and V. S. Bystrov. 1999. Theoretical models of conformational transitions and ion conduction in voltage-dependent ion channels: Bioferroelectricity and superionic conduction. Ferroelectrics 220:157–204.

37. Chapman, D. 1991. Lyotropic mesophases in biological systems. In Liquid Crystals: Applications and Uses, vol. 2. Bahadur, B., editor. World Scientific, Singapore, pp 185–223.

38. Crooker, P. P. 2001. Blue phases. In Chirality in Liquid Crystals. Kitzerow, H-S, Bahr C., editors. Springer, New York, pp 186–222.

39. Leuchtag, H. R. 2008. Voltage Sensitive Ion Channels: Biophysics of Molecular Excitability, Springer, Dordrecht, The Netherlands.

40. Watanabe, A. 1987. Change in optical activity of a lobster nerve associated with excitation. J. Physiol. 389:223–253.

41. Muševic, I., R. Blinc, and B. ŽekŠ. 2000. The Physics of Ferroelectric and Antiferroelectric Liquid Crystals. World Scientific, Singapore.

42. Cohen, L. B., R. D. Keynes, B. Hille. 1968. Light scattering and birefringence changes during nerve activity. Nature 218:438–441.

43. Leuchtag, H. R. 1995. Fit of the dielectric anomaly of squid axon membrane near heat-block temperature to the ferroelectric Curie–Weiss law. Biophys Chem 53:197–205.

44. Palti, Y., and W. J. Adelman Jr. 1969. Measurement of axonal membrane conductances and capacity by means of a varying potential voltage clamp. J. Membr. Biol. 1:431–458.

45. Fishman, H. M, H. R. Leuchtag, and L. E. Moore. 1983. Fluctuation and linear analysis of Na current kinetics in squid axon. Biophys. J. 43:293–307.

46. Wróbel, S., M. Marzak, B. Godlewska, B. Gestblom, S. Hiller, and W. Haase. 1995. Collective and molecular relaxation in ferroelectric liquid crystals. SPIE Proc 2372:169–175.

47. Bystrov, V. S., and H. R. Leuchtag. 1996. Phase transitions in the ferroelectric-active model of ion channels of biomembranes. Ferroelectrics 186:305-307. 1996).

48. Gordon, A., B. E. Vugmeister, H. Rabitz, S. Dorfman, J. Felsteiner, and P. Wyder. 1999. A ferroelectric model for the generation and propagation of an action potential and its magnetic field stimulation. Ferroelectrics 220:291–304.

49. Liu, Y., Y. Zhang, M.J. Chow, Q.N. Chen, and J. Li. 2012. Biological ferroelectricity uncovered in aortic walls by piezoresponse force microscopy. Phys Rev Lett 108:078103.

50. Bystrov, V. S., E. Paramonova, I. Bdikin, S. Kopyl, A. Heredia, C. Pullar, A. L. Kholkin. 2012. BioFerroelectricity: Diphenylalanine peptide nanotubes computational modeling and ferroelectric properties at the nanoscale. Ferroelectrics 440:227–248.

51. Ermolina, I., A. Strinkovski, A. Lewis, and Y. Feldman. 2001. Observation of liquid-crystal-like ferroelectric behavior in a biological membrane. J Phys Chem B 105:2673–2676.

52. Scherowsky, G. 1995. Ferroelectric liquid crystal (FLC) polymers. In: Nalwa, H.S., editor. Ferroelectric Polymers: Chemistry, Physics, and Applications. Marcel Dekker, Inc., New York, pp 435-537.

53. Clark, N.A., and S.T. Lagerwall. 1991. Introduction to Ferroelectric Liquid Crystals. In: Goodby JW et al., editors. Ferroelectric Liquid Crystals: Principles, Properties and Applications. Gordon and Breach, Philadelphia, pp 1–97.

54. Bune, A.V., V.M. Fridkin, S. Ducharme, L.M. Blinov, S.P. Palto, A.V. Sorokin, S.G. Yudin, A. Zlatkin. 1998. Two-dimensional ferroelectric films. Nature 391:874–877.

55. Wada, A. 1976. The α-helix as an electric macro-dipole. Adv. Biophys. 9:1–63.

56. Hol, G.J., P.T. van Duijnen, and H.J.C. Berendsen. 1978. The α-helix dipole and the properties of proteins. Nature 273:443–446.

57. Pikin, S.A. 1991. Structural Transformations in Liquid Crystals. Gordon and Breach, Amsterdam, pp 111–229.

58. Yoshino, K, and T. Sakurai. 1991. Ferroelectric liquid crystals and their chemical and electrical properties. In: Goodby, J.W. et al, editors. Ferroelectric Liquid Crystals: Principles, Properties and Applications. Gordon and Breach, Philadelphia, pp 317–363.

59. Bystrov, V. S., and Leuchtag, H. R. 1994. Bioferroelectricity: Modeling the transitions of the sodium channel. Ferroelectrics 155:19–24.

60. Leuchtag, H. R. 2000. Bioferroelectricity in models of voltage-dependent ion channels. Ferroelectrics 236:23–33.

61. Conley, E. C., and W.J. Brammar. 1999. The Ion Channel Facts Book IV: Voltage-Gated Channels. Academic Press, San Diego.

62. Tytgatt, J., K. Nakazawa, A. Gross, P. Hess. 1993. Pursuing the voltage sensor of a voltage-gated mammalian potassium channel. J Biol Chem 268:23777–23779.

63. Chen, A., C.-D. Poon, T. Dingemans, and E. T. Samulski. 1998. Ferroelectric liquid crystals derived from isoleucine. II. Orientational ordering by carbon-13 separated local field spectroscopy. Liquid Crystals 24:255–262.

64. Lopez, G. A,, Y. N. Jan, and L. Y. Jan. 1991. Hydrophobic substitution mutations in the S4 sequence alter voltage-dependent gating in Shaker K^+^ channels. Neuron 7:327–336.

65. Helluin, O., M. Beyermann, H. R. Leuchtag, and H. Duclohier. 2001. A critical role for the branched sidechain adjacent to the third arginine of the sodium channel voltage sensor. IEEE Trans. Diel. Elect. Insul. 8:637–643.

66. Bendahhou, S., A. O. O’Reilly, and H. Duclohier. 2007. Role of hydrophobic residues in the voltage sensors of the voltage-gated sodium channel. Biochim. Biophys. Acta 1768:1440–1447.

67. Lacrois, J. J., and F. Bezanilla. 2012. Tuning the voltage-sensor motion with a single residue. Biophys. J. 103: L23–L25.

68. Smith-Maxwell, C. J., J. L. Ledwell, and R. W. Aldrich. 1998. Uncharged S4 residues and cooperativity in voltage-dependent potassium channel activation. J. Gen. Physiol. 111:421–439.

69. Posson, D.J., and P.R. Selvin. 2008. Extent of voltage sensor movement during gating of shaker K^+^ channels. Neuron 59:98–109.

70. Logothetis DE, Kammen BF, Lindpaintner K, Bisbas D, Nadal-Ginard B. 1993. Gating charge differences between two voltage-gated K^+^ channels are due to the specific charge content of their respective S4 regions. Neuron 10:1121–1129.

71. Papazian, D.M., L. C. Timpe, Y.N. Jan, L.Y. Jan. 1991. Alteration of voltage-dependence of *Shaker* potassium channel by mutations in the S4 sequence. Nature 349:305–310.

72. Papazian, D.M., and F. Bezanilla. 1997. How does an ion channel sense voltage? News Physiol Sci 12:203–210.

73. Weinreb, G.E., and I.S. Magura. 1998. Physical and molecular basis of ion channel gating: Can electrostatic interactions close the ion channel? Neurophysiology 30:325–327.

74. Zundel, G., B. Brzezinski, and J. Olejnik. 1993. On hydrogen and deuterium bonds as well as on Li^+^, Na^+^ and Be^2+^ bonds: IR continua and cation polarizabilities. J. Mol. Struct. 300:573–592.

75. Kinzel, W. 1983. Directed percolation. In Percolation Structures and Processes. Deutscher, G., R. Zallen, and J. Adler, editors. Adam Hilger, Bristol, and The Israel Physical Society, pp. 425–445.

